# Dynamics of dopamine transients in human olfactory epithelium during reward learning

**DOI:** 10.64898/2026.10.07.757414

**Authors:** L. Paul Sands, Seth R. Batten, Alec E. Hartle, Terry Lohrenz, Jason White, Leonard S. Barbosa, W. Matt Howe, Dan Bang, Peter Dayan, Qiaohan Yang, Adam Dede, Andrew Sheriff, Gregory Lane, Bruce K. Tan, Paul Lenkowski, Christina Zelano, P. Read Montague

## Abstract

Dopamine signaling is crucial for motivation, movement, and learning, but has remained beyond the reach of measurement in healthy humans. Here, we measured sub-second dopamine in the olfactory epithelium of healthy humans during two reward learning tasks with distinct reward types: liquid reward during a conditioning task and monetary reward during an economic exchange game. In both tasks, as trials progress, peak dopamine transients shift from rewarding outcomes back to reward-predictive cues. In the economic game, this pattern retrains quickly when reward magnitude or reward timing is changed without warning. These findings match the time transfer characteristics and amplitude dependence of temporal difference prediction error signals. The results also establish a new, minimally invasive route for recording fast dopamine signaling in healthy humans: the nose.

## Introduction

Artificial reinforcement learning systems now solve wide-ranging problems possessing real world complexity (*1–9*). Biological reinforcement learning systems have long been thought to operate in part through widely projecting monoamine signaling systems (*10–21*). One prominent hypothesis is that dopamine delivery carries a particular style of continuous learning signal that communicates the moment-by-moment difference in expected and received reward: a temporal difference (TD) prediction error (i.e., TD error; *22-24*). A hallmark of dopamine-based TD prediction error signaling is the continuous temporal transfer of dopamine responses across learning from unexpected rewards to reward-predictive cues (*10–12*). Early evidence supporting such a dopamine-encoded TD error derived from action potential recordings in various species (e.g. *10-15* for early work, and *16-21* for later work noting additional heterogeneities in the signals) and BOLD imaging (*25–28*). However, this computational motif has never been tested in healthy humans in experiments possessing chemical specificity and sub-second temporal resolution. Two main obstacles have been the lack of a non-invasive route to record dopamine release directly in healthy humans, and the lack of a method that unambiguously identifies dopamine at sub-second time scales.

In this paper, we address both obstacles by making sub-second monoamine recordings in human olfactory epithelium (*29*) using a validated neural network approach (*30-38*; **Figure 1**). In prior work, we developed a neural network approach to monoamine detection on both carbon and platinum-iridium surfaces (*30-38*; **Figure 1**). These methods have been validated *in-vivo* using transgenic animals where light sensitive proteins can be genetically targeted to specific cell types and optically controlled (*37*; see Methods and **Figure S1**), and where independent fluorophores can also report on dopamine delivery (e.g., dLight; *39*). In separate work (*29*), Zelano and colleagues developed a non-surgical, minimally invasive approach to electrophysiological monitoring of olfactory bulb and olfactory epithelium, which entails positioning FDA-approved stereoelectroencephalography (sEEG) depth sensors (Ad-Tech Macro-Micro; a flexible probe ∼1.3 mm in diameter), equipped with research platinum micro-wire contacts, along the olfactory epithelium just under the cribriform plate (**Figure 1**, **Figure S2**).

**Figure 1.**
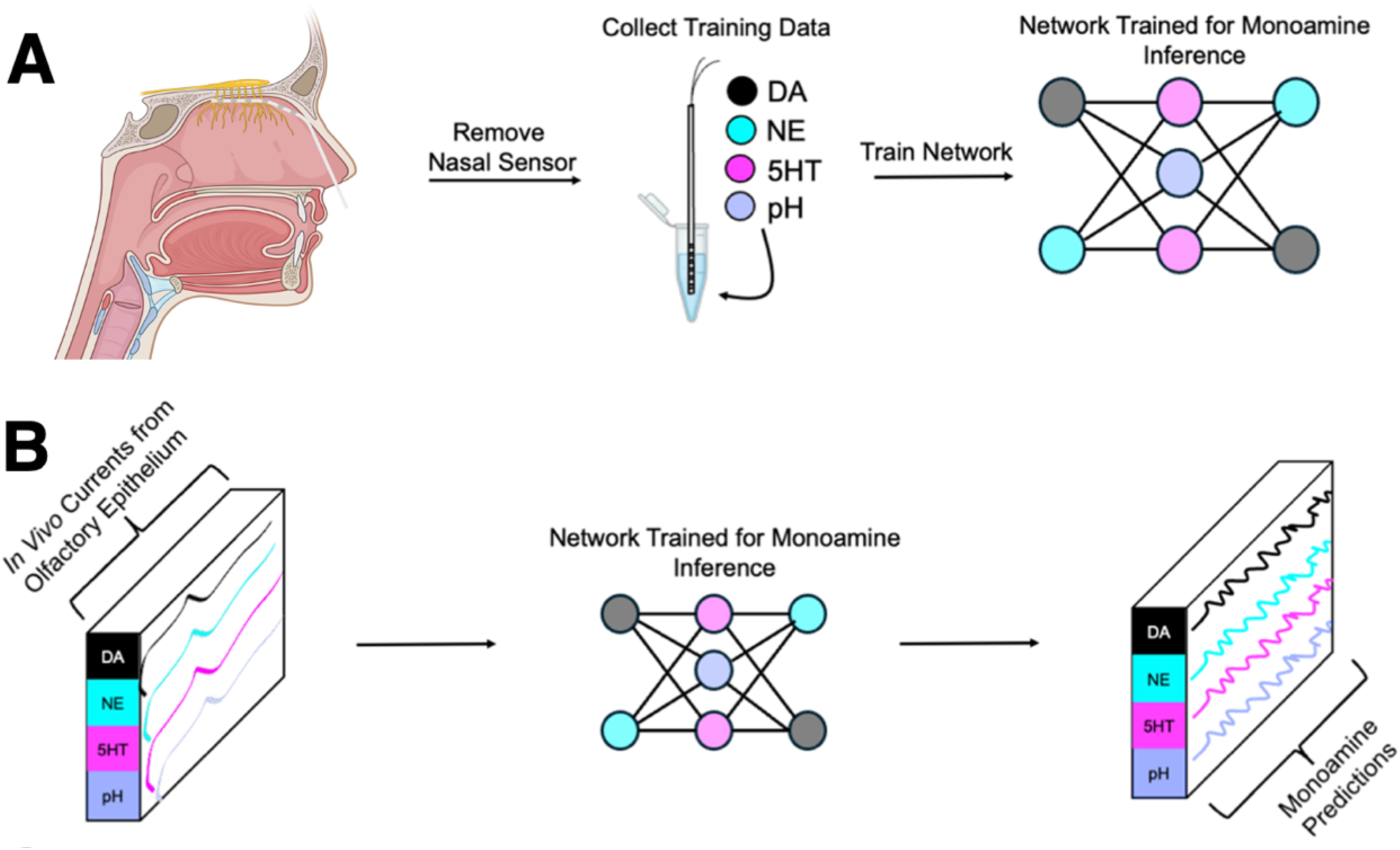
Fast chemical inference in healthy human olfactory epithelium using neural networks. **(A)** Overview of process for recordings from human olfactory epithelium, *in vitro* training data collection, and neural network calibration. Following each experimental recording session (N=46; sensor placement depicted in image at left), the horizontally positioned sensor is removed, and an ensemble of convolutional neural networks is trained from the sensor’s *in vitro* electrical current responses to known concentrations of dopamine (DA), serotonin (5HT), norepinephrine (NE), and pH levels. **(B)** Overview of process for generating vector time series of monoamine concentrations from trained neural network models following *in vitro* model training, testing, and validation. Electrical currents collected *in vivo* from olfactory epithelium are input into the trained network, which output time series concentration estimates for each analyte the model was trained on.

Here, we combined these two approaches by placing sEEG sensors along the olfactory epithelium of healthy volunteers (N = 57) and training neural network models to decode monoamine release in this region during reward learning. In human and rodent olfactory mucosa, the cilia and somata of primary olfactory sensory neurons express dopamine receptor DRD2; furthermore, dopamine release at the level of the olfactory mucosa modulates olfactory behaviors, with genetic or pharmacological blocking of DRD2 activity increasing olfactory sensitivity (*40*). With this nasal recording paradigm in place, we performed five experiments across two task domains that separately use either primary or abstract rewards predicted by cues. The first task, a trace conditioning paradigm with liquid reward, presented a visual pattern (fractal) followed by juice (n = 7) or water (n = 6) for a total of 8 trials (**Figure 2A**). The second task was a well-studied game of monetary exchange, the ultimatum game (*41*), which presents a natural reward prediction interpretation, in which an instruction cue preceeds the onset of a trigger cue that initiates reward harvesting (**Figure 3A-C**). We used the standard ultimatum game (n = 13) as well as two ultimatum game variations where, without warning, we either changed the mean reward magnitude (n = 15) or delayed the timing of the reward (n = 16) to study how monoamine signaling might adapt to changes in task structure.

**Figure 2.**
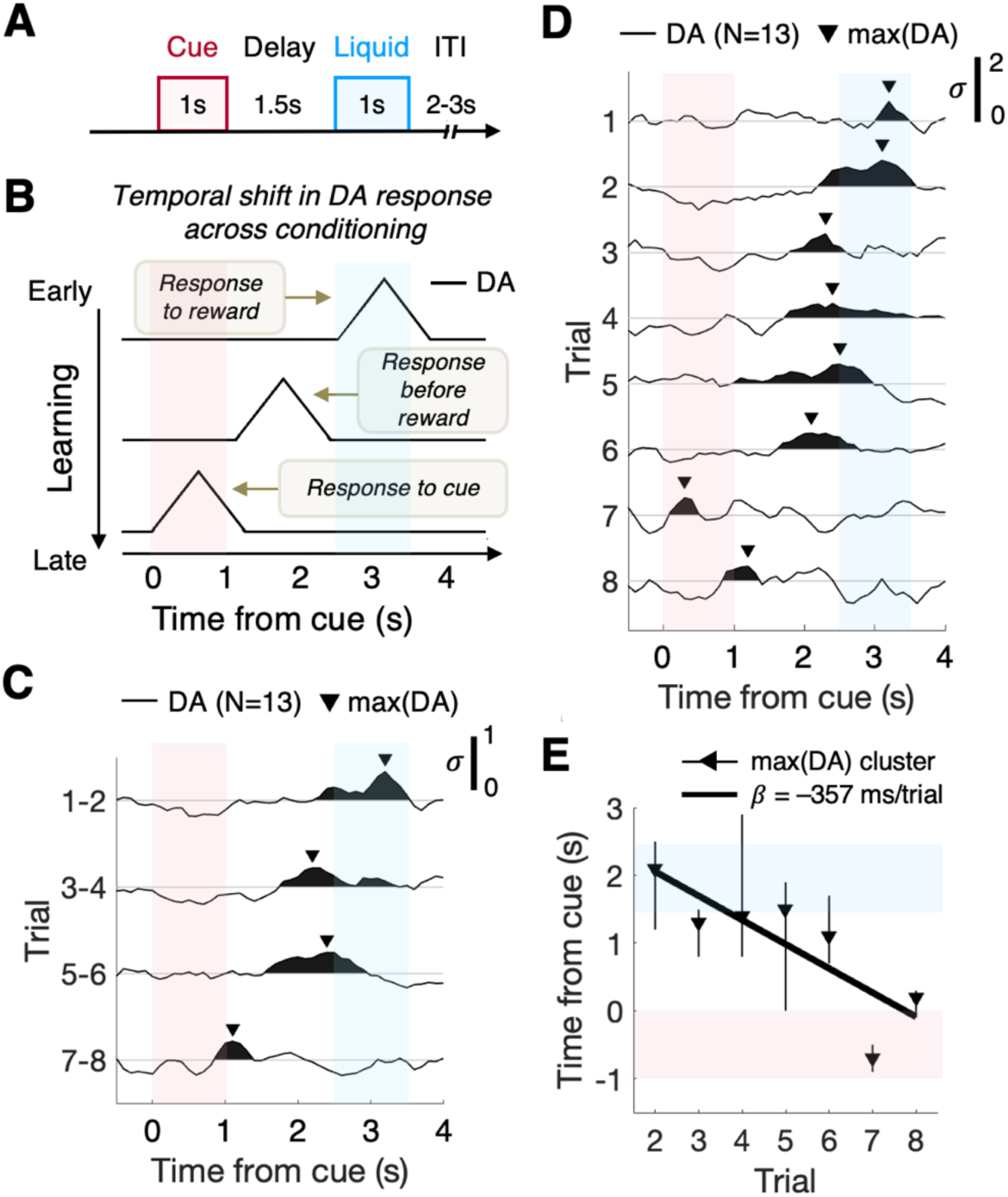
Temporal phase shift of dopamine responses during trace conditioning. **(A)** Liquid trace conditioning task. ITI: inter-trial interval. In all figure panels, red and blue shaded patches indicate the fractal cue period and liquid delivery period, respectively. **(B)** Cartoon depiction of hypothesized temporal shifting in dopamine (DA) fluctuations across conditioning. **(C)** Group-level (N=13) DA time series (z-scored) averaged across participants for blocks of two trials (i.e., block 1 = average of Trial 1 and Trial 2, etc.) and plotted with the peak DA response for each group-level block time series (black triangles), which was identified within a window from 0s to 4s relative to cue onset. Black shaded region beneath DA time series indicates the positive contiguous DA cluster containing the maximum DA level (black triangle). **(D)** Same as **(C)** but plotted for the single-trial DA time series. **(E)** Regression of maximum DA time point on trial number for group-level, single-trial DA time series. Black triangles indicate the timing of the peak DA response; vertical lines represent the onset and offset of the cluster containing the peak DA response (shaded black region in **(D)**). Bold line represents the slope (regression coefficient, *β*) of the group-level DA peak timing shift across trials.

**Figure 3.**
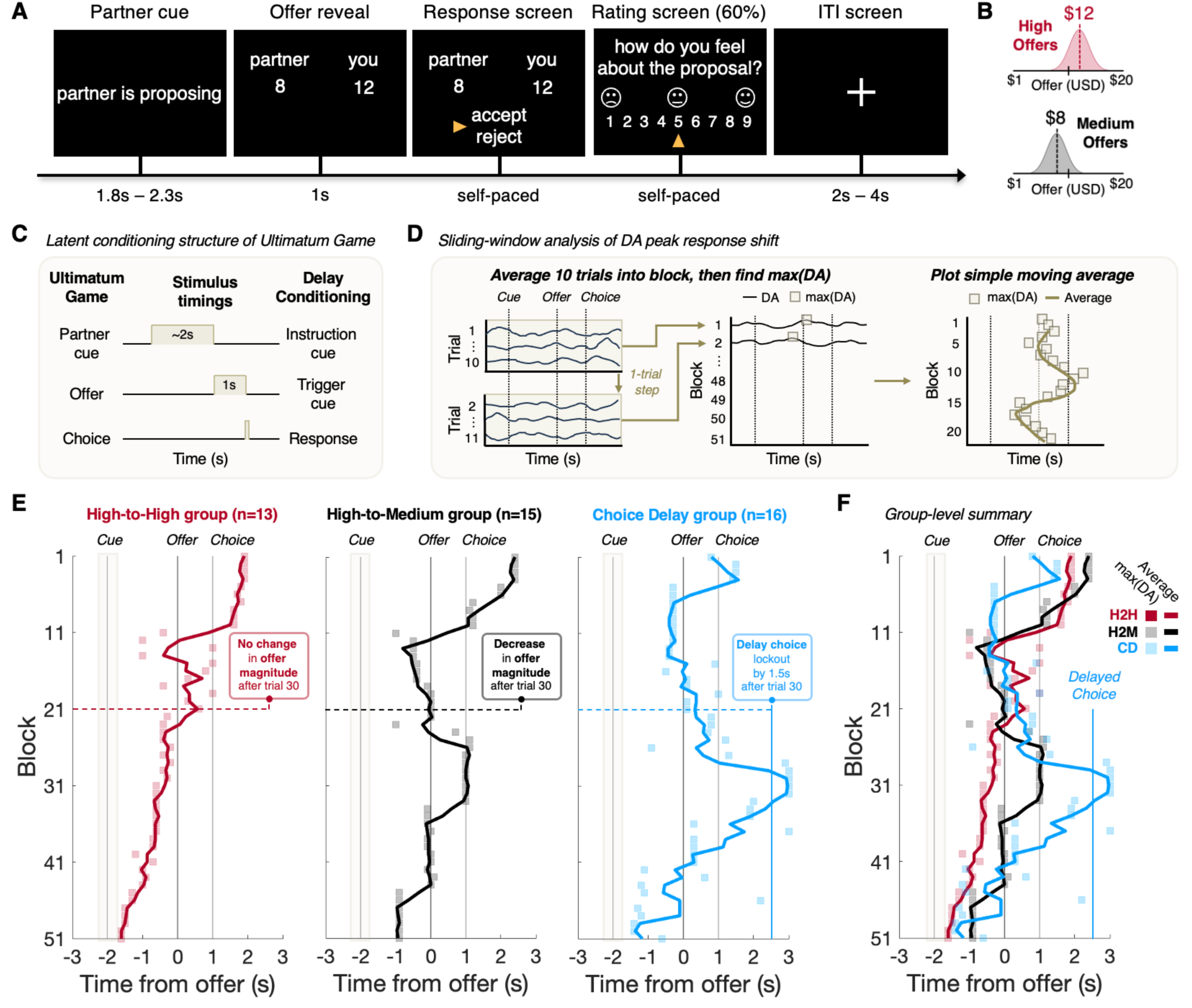
Continuous temporal adaptation of dopamine responses in human olfactory epithelium during two-person economic exchange. **(A)** Example trial of the Ultimatum Game (UG). ITI: inter-trial interval. **(B)** Cartoon depicting high and medium offer distributions. **(C)** Diagram comparing the stimulus timing of UG experiments to the stimulus structure of delay conditioning paradigms. **(D)** Cartoon of sliding-window analysis of dopamine (DA) response shifting within the UG task. Colored squares indicate the DA maximum for each 10-trial block (note the 1-trial step size for sliding window); bold colored line (right) indicates the simple moving average of the DA maximum times over 3 blocks. **(E)** Timing of the maximum DA responses (colored squares) for 10-trial blocks of the UG task for the (left) High-to-High (H2M), (middle) High-to-Medium (H2M), and (right) Choice Delay (CD) cohorts. Bold colored lines indicate the 3-block simple moving average of the peak DA response; vertical lines indicate the timing of trial stimuli; shaded tan region centered on cue onset indicates the jitter in the timing from partner screen to offer reveal. Horizontal dashed colored lines show the transition in offer magnitude or choice timing, or the lack thereof, within the UG task version. **(F)** Group-level results from **(E)** overlaid.

## Results

### Neural network inference of monoamine fluctuations from human olfactory epithelium

Following the completion of the experimental session, we collected *in vitro* calibration data on the sEEG sensor that was placed in the participant’s nasal cavity. We used these data to train a sensor-specific neural network model via cross-validation to predict the concentration of dopamine (DA), serotonin (5HT), and norepinephrine (NE) against varying background pH levels (see Methods; **Figure 1**). The end result of this training process was an ensemble of convolutional neural networks that demonstrated high out-of-sample predictive accuracy for monoamine concentration (**Figure S1B**). Data collected *in vivo* from human olfactory epithelium were then input into the trained sensor-specific model to generate a vector of monoamine time series, which were then aligned to the events of the trace conditioning and ultimatum game tasks.

### Time shifting of dopamine transients in olfactory epithelium during trace conditioning

Thirteen (n = 13) participants completed the liquid trace conditioning task (**Figure 2A**) while dopamine levels were recorded from participants’ olfactory epithelium. On each trial of the task, a square fractal pattern is presented for 1 second before disappearing, leaving a blank screen, and the liquid reward is delivered at a fixed delay of 1.5s following cue offset, with liquid delivery lasting for 1s. The primary liquid reward was either water (n = 6) or apple juice (n = 7), and the same volume of water or juice was given on each trial.

To investigate the adaptations in dopamine responses with learning during trace conditioning (**Figure 2B**), we visualized the average cue-aligned dopamine time series across all participants (n = 13), at the level of single trials and for two-trial averages (**Figure 2C,D**). Further, at the single trial level, we quantified the time point of the maximum positive dopamine level, within a window spanning cue onset to liquid delivery offset (**Figure 2D,E** black triangles indicate maximum). Across aligned trials, we observed a backwards temporal shift of the peak dopamine response within the phase of a trial **(Figure 2C-D**): early in the task, the peak dopamine response was aligned to the reward delivery, and later in the task, the peak response was aligned to cue onset.

We modeled this pattern of adaptation in dopamine responses at the single trial level as a monotonically backwards shift, within the phase of a trial, in the timing of the peak dopamine response (**Figure 2B**), which we tested via linear regression of dopamine peak response time points onto trial number (**Figure 2E**). This linear regression analysis was conducted for the water and juice cohorts separately, as well as for individual participants (**Figure S3**). For each cohort and participant, we performed this analysis on data from trials 2-8 (i.e., the last 7 trials of the experiment). Trial 1 engendered variable dopamine responses to both the fractal cue and liquid reward (**Figure 2D**), though these initial transients are unrelated to experience-based adaptive learning.

For the combined cohort of juice reward and water reward (N = 13), this linear regression analysis confirmed a significant backwards temporal shift in the peak dopamine response as a function of trial (beta = -3.57, *t*(5) = -3.26, *p* = 0.022; **Figure 2E**), with the fitted regression coefficient indicating a per trial adaptation rate in the peak dopamine response of approximately - 350ms/trial. Performing the same linear modeling at the individual subject level, we observed that all 13 participants demonstrated a negative regression coefficient, resulting in a significant negative regression coefficient at the group-level for the combined cohort, as well as for the juice and water cohorts separately (**Figure S3**). Furthermore, we found no difference between water and juice participants’ regression coefficients, indicating equivalent temporal shifting across trials of the peak dopamine responses. The finding of negative regression coefficients across individuals supports the interpretation that the peak dopamine response timing is shifting backwards within the phase of the trial as the experiment progresses.

We replicated the linear modeling analysis above using both the onset and offset of the dopamine response cluster (i.e., contiguous positive dopamine fluctuation) containing the peak dopamine level of each trial (**Figure 2C-E**). These analyses revealed significant backwards temporal shifting across conditioning for these peak response cluster features (**Figure S3**). Lastly, we performed the linear regression modeling of peak response timing and peak response cluster features for serotonin and norepinephrine, which revealed no significant temporal shifting for these other monoamines (**Figure S3**). Thus, the observed patterns in monoamine temporal shifting across conditioning appears selective for dopamine.

### Dopamine signaling in olfactory epithelium during an economic exchange game

A total of forty-four participants (N = 44) completed the ultimatum game (**Figure 3A**). On each trial of the ultimatum game, a proposer (the computer) is endowed with $20 and proposes a split of the money with their partner (the responder; here, the participant), who then decides to accept or reject the proposed monetary split. Importantly, participants are locked out from selecting and submitting their response for 1 second after a trial’s monetary split is first revealed. As such, the offer reveal cues participants that the offered money can be harvested after the 1 second lockout period by performing a button press (accept or reject). Thus, the offer reveal acts as a homolog to a delay conditioning task’s trigger cue (**Figure 3C**).

Each participant performed one of three different versions of the ultimatum game. The first ultimatum game task version, referred to as the High-to-High group (n = 13), represents a control cohort and consists of 60 trials of monetary offers from a partner with a mean of $12 (see Methods; **Figure 3B**). The second task version, referred to as the High-to-Medium group (n = 15), follows the same design as the High-to-High task version, except that the average monetary offer for the final half of the experiment (trials 31-60) switches from the high distribution (mean = $12) to the medium distribution (mean = $8) without warning. Finally, the third task version, referred to as the Choice Delay group (n = 16), follows the same design as the High-to-High task version, with the key difference being an unannounced extension of the response lockout period, from 1s to 2.5s, for the final half of the experiment (trials 31-60). Thus, across all ultimatum game task versions, the first 30 trials were comprised of monetary offers from the high distribution centered on $12, with the response screen onset occurring 1s following the offer reveal.

Similar to our analysis of the trace conditioning data, we investigated the temporal adaptations in dopamine signaling across trials of the ultimatum game. In the ultimatum game, on each trial, the inter-stimulus interval between the partner cue and the offer reveal is jittered, and there is uncertainty in the offer magnitude as defined by the offer distribution. Comparatively, in the trace conditioning task, the timing between cue and reward, and reward magnitude, are kept constant. Given these differences, to facilitate comparison with the observed dopamine signaling dynamics during trace conditioning (**Figure 2**), we sought to focus our ultimatum game analysis on dopamine response patterns that are invariant to these sources of uncertainty in the ultimatum game. To this end, we adopted a sliding-window approach to visualizing the continuous adaptation in dopamine transients across the ultimatum game, using a 10-trial averaging window with 1-trial step size (**Figure 3D**). Specifically, for each participant, we first averaged dopamine time series across blocks of 10 trials (i.e., block 1 = trials 1-10, block 2 = trials 2-11, etc.), and then averaged the block-level dopamine time series data across all participants within each cohort. From these cohort-level datasets of block-level dopamine time series, we identified the timing of the peak dopamine response level within a window spanning offer reveal and response screen onset (see Methods). We additionally performed this analysis using contiguous, non-overlapping 10-trial blocks (**Figure S4**), as well as for sliding-windows of different trial widths (**Figure S5**), which confirmed the robustness of this approach.

As shown in **Figure 3E**, for all three ultimatum game cohorts, we found that the timing of the peak dopamine response moved backwards within the phase of a trial from the reward time (i.e., response selection) to align at or before the trigger cue (i.e., offer reveal) as the experiment progressed through the first 30 trials (**Figure S3**). Across the final 30 trials of the ultimatum game, for the High-to-High cohort, the timing of the peak dopamine response continued to shift backwards gradually, from before the trigger cue to the partner cue onset (**Figure 3E**, left). For the High-to-Medium group, the peak dopamine response timing first re-aligned back to the reward time (response selection) and then repeated the backwards temporal shift across the remainder of the experiment to align to before the trigger cue (**Figure 3E**, middle). For the Choice Delay cohort, we observed a similar re-tracing of the peak dopamine response timing displayed over the first 30 trials, with the crucial exception being that the timing of the peak dopamine response first aligned to the newly delayed reward time at 2.5s post-offer reveal–overshooting the initially-learned reward time (i.e., at 1 second post-offer)–before reversing and shifting backwards within the phase of a trial to align to before the onset of the trigger cue (**Figure 3E**, right). We found that these patterns of temporal shifting in dopamine responses for each cohort were consistent using different levels of trial averaging (**Figure S5**) and were observable at the single-subject level (**Figure S6**). Furthermore, linear modeling of the observed dopamine response dynamics across ultimatum game cohorts revealed statistically significant patterns in dopamine temporal adaptations that were not observed for serotonin or norepinephrine (**Figure S4**). In all, these patterns match what one could expect from a continuous TD error signal encoded by dopamine fluctuations in olfactory epithelium.

## Discussion

In this report, we developed and deployed a method for recording sub-second monoamine signaling from the olfactory epithelium of healthy human subjects in order to eavesdrop specifically on fluctuating dopamine delivery during two reward learning tasks. The first task comprised a trace conditioning task, where a visual cue predicted the near-term delivery of apple juice or water. This simple experiment revealed multiple features characteristic of a TD prediction error in the observed dopamine dynamics: dopamine fluctuates positively when liquid reward is unexpectedly delivered, and this response monotonically shifts from the time of the reward to the time of the cue as the 8 trials progress (**Figure 2**). Crucially, we note that the trials of the trace conditioning task (and ultimatum game) are significantly staggered in time; thus, these observed changes in the timing of dopamine responses are relative to the phase of the trial (i.e., the temporal structure of trial stimuli), and are not attributable to oscillatory or random fluctuations in dopamine levels across the experiment. Additionally, participants’ breathing rate was restful and consistent throughout task performance, and given the temporal jittering of trials within each task, the observed dopamine effects are not due to cue-timed breathing responses. These observations argue for a dopamine-based, TD-like prediction error signal using primary reward, but available at the level of healthy human olfactory epithelium. Although the trace conditioning task consists of a relatively small number of trials, the measured dopamine fluctuations track well with the kind of learning rates derived from spike rate changes in primate dopamine neurons when reward timing is changed suddenly (*13*). However, in rodents such paradigms show a much slower continual shift, taking hundreds of trials for dopamine responses to shift back to predictive cue (*42*).

We found that the same basic pattern of dopamine signaling during trace conditioning was also observed during the ultimatum game across the first 30 trials for all three subject groups (**Figure 3**): the maximum dopamine fluctuations initially occurred in response to the reward and with further trials moved backwards within the phase of the trial towards the trigger cue (offer reveal). In the ultimatum game, additional strong evidence for a TD-like dopamine signal emerged from the effects of task manipulations, where one of two sudden changes to task structure, decreasing mean offer magnitude or delaying choice timing, induced a shift in the relative phase of the peak dopamine response. For the High-to-Medium group, a sudden shift at trial 31 in the reward distribution, from high offers to medium offers, correlated with a forward shift of the dopamine maximum to just after the response screen onset; with repeated trials drawing from the medium distribution (while holding response timing constant), the dopamine maximum again shifted backwards within the phase of the trial to occur in response to or before the offer reveal (**Figure 3E**, middle), in a fashion consistent with a TD-like signal. Comparatively, for the Choice Delay cohort (**Figure 3E**, right), while the reward distribution was held constant over the last 30 trials of the task, the response lockout duration was increased from 1 second to 2.5 seconds. Remarkably, this change correlated with a forward shift in the peak dopamine response time, migrating from the offer reveal to the new response screen onset time (i.e., at 2.5s post-offer); with repeated trials, the peak dopamine timing reversed and progressed backwards to align to before the proposal screen.

Collectively, the present data provide strong positive evidence for a TD-like prediction error encoded by dopamine (and not norepinephrine or serotonin; **Figure S3,S4**) at the level of the olfactory epithelium. Across multiple tasks engaging adaptive learning from primary and abstract rewards, the observed olfactory epithelium dopamine dynamics resembled a dopamine signal that would be expected on the basis of recordings from dopamine neurons in primate midbrain (*13*) and the connection to TD learning models (*11,12*). The similarities in observed dopamine dynamics across trace conditioning, a task directly engaging retro-nasal olfaction through tasting of liquid rewards, and the ultimatum game, a non-olfactory task of social economic decision-making, suggest that the observed dopamine signaling patterns are global, are not context-dependent, and are accessible at the peripheral olfactory system. Indeed, for a crucial broadcast signaling system like mesencephalic dopamine, it is possible that a substantial branch of this system’s information may be available at the level of the olfactory epithelium to modulate olfaction, in addition to known local sources of dopamine release to this site (e.g., superior cervical ganglion; *40*). While alternative explanations remain to be investigated, such as a never-before-described collection of dopamine neurons that project to olfactory epithelium, the present data suggest a connection of midbrain dopamine signaling to one of the evolutionarily oldest sensory systems, chemical sensing. Additional work will be required to support this new window on the human dopamine system, but this new, minimally invasive method for monitoring fast dopamine signaling in healthy humans could substantially facilitate our investigation of this vital neuromodulatory system and its breakdown in disease and injury.

## Acknowledgements

We thank the volunteers who participated in this study and without whom this research would not have been possible.

## Author contributions

Conceptualization: L.P.S., S.R.B., T.L., P.D., C.Z., P.R.M.; Data curation: L.P.S., S.R.B., A.H., J.W., L.S.B., Q.Y., G.L., C.Z.; Formal analysis: L.P.S., S.R.B.; Funding acquisition: T.L., G.L., C.Z., P.R.M.; Investigation: L.P.S., S.R.B., A.H., Q.Y., A.D., A.S., G.L., C.Z.; Methodology: L.P.S., S.R.B., A.H., T.L., J.W., L.S.B., W.M.H., Q.Y., A.D., A.S., G.L., B.T., P.L., C.Z., P.R.M.; Project administration: T.L., G.L., B.T., P.L., C.Z., P.R.M.; Resources: B.T., P.L., C.Z., P.R.M.; Software: L.P.S., S.R.B., T.L., J.W., L.S.B.; Supervision: T.L., G.L., B.T., P.L., C.Z., P.R.M.; Validation: L.P.S., S.R.B., A.H., T.L., J.W., L.S.B., W.M.H., D.B., P.D., Q.Y., A.D., A.S., G.L., B.T., P.L., C.Z., P.R.M.; Visualization: L.P.S., S.R.B., P.R.M.; Writing – original draft: L.P.S., S.R.B., P.R.M.; Writing – review & editing: L.P.S., S.R.B., A.H., T.L., J.W., L.S.B., W.M.H., D.B., P.D., Q.Y., A.D., A.S., G.L., B.T., P.L., C.Z., P.R.M.

## Data, Code, and Materials Availability

All data and code needed to evaluate and reproduce the results in the paper are present in the paper and/or the Supplementary Materials. The neurochemical time series data included in this manuscript, and accompanying code for recreating main text figures, can be found at **<u>ZENODO link upon acceptance</u>**. This study did not generate new materials.

## SUPPLEMENTARY MATERIALS

### Materials and Methods

#### Participants

##### Human participants and ethics

This study included N = 57 datasets from 46 recording sessions across 42 individual healthy participants (28 female, mean age 36 ± 13 (SD), range: 19-75 years; **Table S1**). At each recording session, a participant had a unique sensor placed in the nasal cavity at the level of the olfactory epithelium (**Figure 1A**, **Figure S2**), and monoamine levels were measured during behavioral tasks. Across all participants, 46 nasal cavity recording sessions were completed (each session was comprised of one or more experimental task recordings), resulting in a total of 46 unique sensors included in the calibration panels of **Figure 1C**. A total of 12 individual participants completed the trace conditioning task, with one participant completing both the Water and Juice trace conditioning task versions on two different recording session visits, resulting in a total of n = 13 trace conditioning datasets. A total of 40 individual participants completed the ultimatum game experiments, with 4 participants completing two different ultimatum game versions on two different recording session visits, resulting in a total of n = 44 ultimatum game datasets. Note that 11 participants completed both the trace conditioning and ultimatum game experiments. Written and verbal consent were obtained from all participants before being included in the study. This study complied with all ethical regulations and was approved by the Institutional Review Board of Viriginia Tech (protocol #25-359) and Northwestern University (STU00218720).

##### Rodent models and ethics

DAT-Cre mice (N=2; **Figure S1C**) were used for optogenetic experiments. The mice were injected with an Adeno-associated virus (AAV) to drive Cre-recombinase-dependent expression of channelrhodopsin (ChR2, AAV-DIO-hChR2(H134R)-EYFP, Addgene viral prep #26973-AAV9) in midbrain dopamine neurons. Following viral injection and optic fiber surgery (see below), mice were housed individually. All rodent experiments were conducted in accordance with the ethical guidelines of the National Institutes of Health and were approved by the Institutional Animal Care and Use Committee at Virginia Tech, Blacksburg, VA (protocol #23-112).

#### Behavioral testing

##### Trace conditioning task

A subset of participants (n = 13) performed a trace conditioning task in which a fractal visual cue is shown for 1 second before disappearing and is followed by the delivery of either apple juice (n = 7) or water (n = 6) at 1.5 seconds post-cue offset (**Figure 2A**). The task consisted of 8 conditioning trials, with each trial separated by a jittered 2-3 second inter-trial interval (ITI). Liquid was delivered via a Harvard Apparatus pump (Model 33, Catalog # 55-3333) containing a 50 mL syringe of liquid connected to a Tygon tube (B-44-3, AAB00004; ID: 2.4 mm, OD: 4 mm) that was placed in the mouth of the participants. The participants were instructed to simply watch the screen, and liquid was passively delivered to the participants via the pump after cue offset (∼0.8 mL total for each delivery; this amount just covers the tongue). The pump was placed outside of the experimental room to minimize participants detecting pump noise during liquid delivery. The game was controlled by custom written software using HTML and JavaScript and was displayed on a desktop monitor.

##### Ultimatum game

A subset of participants (n = 44) played the ultimatum game. In this task, all participants played as the responder and the computer acted as the proposer. The participants were presented with 60 trials of offers from the computer proposer. The participants were instructed that they must accept or reject the offer on each trial, and that if the participants accepted the offer, then both parties received the proposed money, but if they rejected the offer, then both parties got nothing. On 60% of the trials the participants were asked to rate how they felt about the previously proposed offer on a 1-9 scale. Participants were instructed that they would receive the monetary outcome of one random trial from the task at the end of the experiment as a monetary bonus compensation for task completion.

On each trial of the UG task (**Figure 3A**), participants were presented a screen stating “Partner proposing” for 1.8-2.3 seconds (jittered) followed by an offer reveal screen displaying the trial’s offer for 1 second. As such, the participants are locked out from making a decision for 1 second, after which a selector arrow is displayed on the screen allowing participants to make their decision (self-paced). Each trial was separated by an inter-trial interval (ITI) of 2-4 seconds. The game was controlled by custom written software using HTML and JavaScript and was displayed on a desktop monitor. Participants made their decisions using two button boxes, one placed in each hand.

Participants performed one of three different UG task versions. In the High-to-High group (n = 13), the offers for trials 1-60 were drawn from a high distribution (mean = $12, SD = $1.5); the participants were not notified of the offer distribution or expected offer magnitudes. In the High-to-Medium group (n = 15), offers for trials 1-30 were drawn from a high distribution (mean = $12, SD = $1.5), whereas offers for trials 31-60 were drawn from a medium distribution (mean = $8, SD = $1.5); the participants were not given any information regarding the offer distributions or that they would switch during the task. Lastly, in the Choice Delay group (n = 16), offers for trials 1-60 were drawn from the high distribution (mean = $12, SD = $1.5), but for trials 1-30, participants were locked out from making a decision for 1 second, whereas for trials 31-60, participants were locked out from making a decision for 2.5 seconds; the participants were not instructed regarding the offer distribution or the duration of the choice lockout period across trials.

#### Nasal cavity sensor

##### Sensor preparation

We used an AD-Tech Macro-Micro sEEG depth sensor (MM16A-SP05X-000) for all monoamine recordings from olfactory epithelium. This sensor is a 1.3 mm diameter polyurethane tube that includes 6 platinum-iridium macro-wire contacts spaced 5 mm apart center-to-center. Platinum-iridium micro-wire contacts (38 µm in diameter) are positioned between the distal 4 macro contacts (spaced 5 mm center-to-center) such that there are 3 micro contacts positioned around the polyurethane tube in front of the first macro contact, 3 micro contacts positioned around the polyurethane tube in front of the second macro contact, 2 micro contacts positioned around the polyurethane tube in front of the third macro contact, and 2 micro contacts positioned around the polyurethane tube in front of the fourth macro contact (**Figure S1,S2**). A small nasal sponge (Merocel) was placed on the sensor such that a row of 4 micro contacts would face up toward the olfactory epithelium and the cribriform plate upon nasal cavity placement. The sponge was used to help stabilize the sensor and to ensure the correct directionality of the sensor during placement. Measurement marks were placed on the surface of the polyurethane tube, registered to each participant’s nare entry, to allow for additional precision during placement and to assess if the sensor moved during the experiment. Both sponge placement and nare measurements were participant specific; thus, each sensor was modified on a per person basis.

##### Sensor placement

Sensors were placed based on computed tomography (CT) scan imaging and measurements of individual nasal anatomy. An experienced ENT (coauthor’s P.L. and B.K.T.) placed the sensors under endoscopic and navigational guidance along the superior surface of the nasal cavity, just below the cribriform plate. When placed, the sensor sits inside the nasal cavity on the olfactory epithelium, ∼1 mm away from the olfactory bulb. In preparation for sensor placement, the majority of participants were given a 1:1 4% lidocaine and Afrin spray during the decongestant phase followed by the placement of pledgets soaked in a solution containing a 4:1 mixture of 4% lidocaine and 1/1000 epinephrine. Some participants chose to solely receive decongestant (1% solution phenylephrine or Afrin). Decongestant is used to open the anterior nasal cavity before sensor placement but is not applied to the olfactory cleft. Sensor placement involves only brief minimal discomfort that resolves immediately when the sensor is in place; however, some participants still prefer topical anesthetic. Proper sensor positioning was confirmed by a post-placement CT scan (**Figure S2**).

##### CT scan

Pre-placement CT scans for each participant were acquired approximately 1 week before a participant’s study session so that nasal anatomy could be assessed and measurements could be taken for sensor modifications. This pre-placement CT also aided as a screen to make sure that a participant’s anatomy was amenable to sensor placement. CT scans were acquired with a cone beam CT machine (Xoran) with low-dose (participant effective dose estimate = 0.032-0.164 mSv) sinus-scan protocol. The sensor modifications, placement, and CT scanning procedure were adapted from (*29*).

#### Human monoamine measurements

##### Overview

We performed monoamine measurements using our chemical inference approach driven by artificial intelligence (*34, 36–38*). Specifically, we apply a small voltage ramp to the nasal sensor micro contacts and measure back the resulting electrical current in the nasal cavity. These electrical currents carry information regarding the chemical species present and their concentrations. After *in vivo* nasal cavity data acquisition, we remove the sensor from the nasal cavity and collect *in vitro* training data that is used to train an AI model (i.e., an ensemble of convolutional neural networks (CNN); *34, 36-38*) to predict monoamine concentration from the electrical currents collected in the nasal cavity. Specifically, we expose the sensor to known concentrations of dopamine, serotonin, and norepinephrine against a background of varying pH while applying the same voltage ramp and measuring back the electrical current response. We then use the monoamine labels, concentration labels, and the *in vitro* electrical currents collected in the controlled wet lab environment to train an AI model via out-of-sample cross-validation (**Figure 1A**); this model can then generate time series estimates for the analytes it was trained on. After training and validating the AI model is completed, the *in vivo* electrical currents collected in the nasal cavity are used as input to the trained model, which in turn generates time series of the predicted monoamine fluctuations within the nasal cavity during the experimental recording session (**Figure 1B**).

##### Nasal cavity electrochemical data acquisition protocol

To measure monoamine levels in human nasal cavity, we applied a triangular voltage ramp (hold at -0.6 V for 90 ms, ramp up from -0.6 V to +0.6 V at 240 V/s, ramp down from +0.6 V to -0.6 V at -240 V/s) at 10 Hz to the micro contacts on the sensor, using one micro contact as a working surface and another as a reference surface, and measured the resulting current response, which carries information about the chemical milieu in the nasal cavity. We collected data for 1-2 min before starting any behavioral tasks to allow for electrode equilibration. The recordings were collected using Molecular Devices (Molecular Devices, LLC, San Jose, CA, 95134) equipment: the Axon Instruments MultiClamp 700B Patch Clamp Amplifier, the Electrochemistry Headstage (CV 7B-EC; slightly modified to allow larger current measures), and the Digidata 1550B Data Acquisition System.

##### In vitro data acquisition for training the AI monoamine models

We built a sensor-specific monoamine model for each sensor used in this study. For each sensor, four *in vitro* datasets were collected, one for each analyte (dopamine, serotonin, norepinephrine, and pH). The *in vitro* data acquisition equipment was of the same type (same component type/number) used in the *in vivo* nasal cavity data collection. For each of the three monoamine datasets, we collected 30 distinct concentrations of the analytes dispersed at equal intervals over the range from 0 to 2500 nanomolar (nM), with a pH of around 7.4, while the other two analytes were kept at 0 nM. In addition, each of these datasets contained 5 mixture solutions at 840 or 1680 nM, where more than one monoamine was present in the calibration solution. For the pH dataset for each sensor, we collected 11 pH values in the range 7.0-7.8, with the concentration of the three monoamines was set to 0 nM; we also collected 5 mixture solutions in the pH dataset, as described above. The concentrations were randomized within each monoamine, as was the order in which analytes were collected per sensor.

To collect the *in vitro* calibration data, each sensor was placed vertically into a 1.5 mL Eppendorf tube containing the monoamine concentration of interest, such that the working and reference micro-wire contacts were submerged. We then recorded electrical current responses at 10 Hz for 65 seconds using the same triangular voltage measurement protocol deployed in the nasal cavity. For each concentration, we selected the most stable continuous 15-second section (i.e., 150 current responses) from the final half of this 65 second window, which was chosen to reduce variation between electrical current responses for each concentration due to electrical noise and sensor equilibration.

##### Training AI model for monoamine concentration prediction

For the training of each sensor-specific AI monoamine model, we adopted a mixture-of-experts approach, where the average of an ensemble of 5 training runs was used to generate the final monoamine concentration predictions. Each single training run involved a calibration dataset containing 150 differentiated electrical current sweeps from each unique analyte concentration. Each calibration dataset was split into a training set, containing 90% of the data, and a validation set containing the remaining 10%. Each calibration dataset was also split by concentration, such that all data points of any given analyte concentration were included in the same training set or validation set. To avoid zero gradients during each training run, we normalized the calibration dataset by z-scoring within analyte and then shifting by 10 standard deviations for each analyte. After training, the predictions from each training run were subjected to the inverse of this normalization procedure. The models were trained for 100 epochs.

##### Evaluating the trained monoamine models

A 10-fold cross-validation test was performed for the monoamine model per sensor. The training data were split into 10 folds without overlapping entries. Each fold was held out as a test set, and an ensemble of models were created via a training run on the remaining data. Ensembles of four models were used instead of five to reduce computation time. The mean of the predictions on the test set from each 4-model ensemble was calculated for each fold, resulting in predictions of concentrations from ensembles of models that had never been exposed those concentrations. The resulting predictions from each fold were used to compute the root mean squared error (RMSE) per analyte, and further binned and averaged for display and to compute the r-squared statistic (R^2^). See **Figure S1B** for the results of this procedure.

##### Generating in vivo monoamine estimates from the nasal cavity using trained AI models

We generated nasal cavity monoamine estimates using the same ensembled model described above. Electrical currents from the nasal cavity were differentiated as in the training data and then input to each ensemble model, and the averaged predictions across the ensemble of models (mixture of experts) were used as the final monoamine concentration estimates.

##### Monoamine model architecture

We modified the InceptionTime time series classification model (*43*) to perform multi-variate regression. The model was coded in Python using TensorFlow (*44*) and Keras (*45*). Specifically, the modified InceptionTime network is based on two ResNet (*46*) blocks. Each block contains 3 convolutional blocks, with a convolutional block composed of 4 convolutional layers in parallel, each with 32 filters and with increasing kernel sizes (1, 10, 20 and 40). The output of each of these convolutional layers is stacked together, after which batch normalization and rectified linear unit (RELU) activations are applied. This output passes through a bottleneck convolutional layer with kernel size 1 and 32 filters; the output of this bottleneck layer serves as the input for the next convolutional block. To start the implementation of the ResNet architecture, the input data, after passing through a bottleneck convolutional layer with kernel size 1 and 32 filters, is added to the output of the first ResNet block, and, after activation, this serves as the input to the second ResNet block. The output of the second ResNet block is passed through a global average pooling layer and then through a dense layer with four output nodes, which yield the predictions for each of the four analytes. All activation functions are RELU, except for the final dense layer, which uses softplus activation.

All models were evaluated using the mean squared error loss. Model training used the ADAM optimizer (*47*), with an initial learning rate of 1e^−3^ that halved after 5 successive epochs without a decrease in validation loss. The models had a minimum learning rate of 1e^−6^, and the batch size was 64. The loss on the validation set was calculated after each epoch. The model from the epoch with the lowest validation loss was selected as the final model for that training run.

We generated the final predictions for nasal cavity monoamine levels using a mixture of experts, where we train an ensemble of models with the same hyperparameters and average their predictions. This ensembling approach accounts for differences in the weights generated during initialization, variations in the order in which data is fed to the algorithm during training, and the stochastic descent algorithm, which all introduce variation to the convergence of each model. Additionally, different validation sets were used in each run to avoid overfitting. See (*34*) for previously published methods.

#### Rodent monoamine measurements and optical stimulation

##### Overview of rodent optogenetic experiments for in vivo validation

In addition to demonstrating the *in vitro* validation of the monoamine AI model trained on Ad-Tech Macro-Micro sEEG depth sensors used in the human nasal recordings (**Figure S1A,B**), we extended this validation to the *in vivo* setting by leveraging causal optogenetic stimulation of rodent dopamine neurons to measure changes in dopamine levels in response to active and null optical stimulation (**Figure S1C-E**). In our prior work using carbon fiber-based sensors, we used optogenetic excitation of dopamine, serotonin, and norepinephrine neurons in mice to demonstrate the *in vivo* validity of the monoamine AI model training protocols (described above) and the specificity of changes in monoamine levels in response to selective optical excitation of the parent neural populations and in response to active but not inactive optical stimulation (*37*). Given that the large diameter of the Ad-Tech Macro-Micro sEEG depth sensors (1.3mm) prohibits its deployment in rodent neural tissue, we elected to perform the *in vivo* validation experiment using the Ad-Tech Behnke-Fried depth sensor (**Figure S1A**) that possesses a bundle of platinum micro-wire contacts that are of equivalent material and comparable surface area as the Macro-Micro depth sensor’s platinum micro-wire contacts (see **Figure S1A** for cartoon comparison). To this end, we first trained a sensor-specific monoamine AI model for Behnke-Fried sensors *in vitro* to predict out-of-sample concentrations of dopamine against varying levels of pH, following the same process described above for the Macro-Micro depth sensors. Note that this sensor-specific monoamine AI model trained for the Behnke-Fried sensor did not include serotonin or norepinephrine *in vitro* training data; thus, the present rodent experiments using optogenetics specifically address the *in vivo* validation of the dopamine concentration predictions from platinum surfaces. Following the *in vitro* monoamine AI model training for the Behnke-Fried sensor, we deployed this sensor to measure dopamine release in the striatum while selectively optically stimulating (with active or null optical excitation) dopamine neurons in the substantia nigra pars compacta (**Figure S1C**). Following completion of the optogenetic experiments, the *in vivo* electrical currents collected in the dorsal striatum are used as input to the sensor-specific monoamine AI model, which in turn generates time series of the predicted dopamine fluctuations in response to optical stimulation conditions (**Figure S1C-E**).

#### In vivo rodent data acquisition

We injected DAT-Cre mice (N=2) with an Adeno-associated virus (AAV) to drive Cre-recombinase-dependent expression of channelrhodopsin (ChR2, AAV-DIO-hChR2(H134R)-EYFP, Addgene viral prep #26973-AAV9) in midbrain dopaminergic cells (from bregma, AP: -3.16mm, ML: +/- 1.35mm, DV: -4.2mm), following our previously published work (*37*). We positioned an optic fiber (400 *μ*m, Doric) at a distance of 75 *μ*m above the site of viral infusion to enable light delivery to activate the opsin. For recordings of optogenetically evoked dopamine release, mice were anesthetized using isoflurane (1-2%) delivered via a SomnoSuite system (Kent Scientific; Torrington, CT) and were placed in a stereotaxic frame (Kopf Instruments; Tujunga, CA). The Behnke-Fried electrode was then implanted into the dorsal striatum; this implantation occurred approximately 4-6 weeks after AAV infusion.

To measure dopamine levels in rodent dorsal striatum, we inserted the Behnke-Fried electrode into the dorsal striatum (AP: 0.75mm to 0.80mm, ML: +/- 1.45mm, DV: -3.05mm to -3.20mm) and applied a triangular voltage ramp (hold at -0.6 V for 90 ms, ramp up from -0.6 V to +0.175 V at 155 V/s, ramp down from +0.175 V to -0.6 V at -155 V/s) at 10 Hz to the micro-wire contacts on the sensor, using one micro-wire contact as a working surface and another as a reference surface. Note that the shape of the forcing function of this electrochemistry acquisition protocol for the Behnke-Fried electrode was the same standard triangular waveform as used for the Macro-Micro depth sensors used in the human nasal recordings, although with a smaller peak voltage. We used this electrochemical recording protocol and the *in vitro* calibrated monoamine AI model for the Behnke-Fried electrode to produce time series estimates of dopamine concentration in response to optogenetic stimulation (**Figure S1C**). We separated stimulation evens by approximately 3 min, which was controlled using a manual trigger on a waveform generator (AFG1062, Tekronix) and time-locked by a TTL. To control for non-specific effects of optical stimulation, we inter-leaved trials using light delivery with the same waveform parameters but using light red-shifted outside the excitation wavelength of ChR2 (617 nM LED, Thorlabs). We used 3 stimulations with active and inactive optical stimulation to generate results shown in **Figure S1D** for an individual mouse, and we used all 6 trials of active and inactive optical stimulation for averaging across mice in **Figure S1E**.

#### Breathing data acquisition

We collected breathing data throughout each experimental recording by placing a breathing cannula (Sniff Logic) in the nostrils of the participants connected to the BiNAPS Diagnostic Pressure Transducer (AirLife) machine. The BiNAPS was connected directly to the electrochemical data collection hardware, such that the nasal cavity electrical currents carrying monoamine information and the voltage changes from breathing were collected by the same hardware and software.

#### Data analysis

##### Monoamine data pre-processing

Monoamine time series data for the human trace conditioning task and the ultimatum game task, as well as the rodent optogenetic experiments, were aligned to the experimental time course using MATLAB (2024b, Mathworks, Natick, MA). For each of the 8 trials of the trace conditioning task, we segmented each monoamine time series into 7-second snippets aligned to the offset of the fractal cue, from 3 seconds before fractal cue offset to 4 seconds after cue offset. For each of the 60 trials of the ultimatum game, monoamine time series were segmented into 6-second snippets centered on the onset of the offer reveal screen (i.e., 3 seconds before and after offer reveal). The resulting trial-level monoamine time series for each participant and for each task were linearly detrended, causally smoothed with a 0.5 second window (i.e., 5-sample window), and then normalized via z-scoring (i.e., each trial is z-scored independently using all data within the trial snippet). The rodent monoamine time series data were processed in the same manner as the human monoamine time series data, except that a 1 second window (i.e., 10-sample window) was used for causal smoothing, and the data were normalized at the trial level using z-scoring based on the mean and standard deviation of a 0.5 second pre-stimulation baseline period (**Figure S1D,E**). All subsequent analyses were performed on the normalized, trial-level monoamine time series.

##### Rodent neurochemical data analysis

For the rodent optogenetic experiments, to determine the mean dopamine response to active and inactive optical stimulation, we took the average of 3 trials of active stimulations and 3 trials of inactive (null) stimulations (**Figure S1D**) and visualized the differences between conditions for an individual mouse (**Figure S1D**) and for both mice’s data averaged together (**Figure S1E**). Note that each mouse contributed three active optical stimulation trials and three inactive optical stimulation trials (i.e., n = 6 trials per condition in **Figure S1E**).

##### Trace conditioning data analysis and linear modeling

We investigated adaptations in the timing of monoamine responses across trace conditioning by quantifying the timing of the peak monoamine response level on each trial of the task and linearly regressing these response time points against trial number. We performed this analysis for each participant individually (**Figure S3B**, bottom), as well as for group-level averaged monoamine time series data, in which the participants of a given cohort were first all averaged together (yielding a single, 8-trial dataset for each monoamine) before performing the peak response timing regression analysis. This group-level analysis was performed separately for the Water cohort and Juice cohorts (**Figure S3B**, top), and for the combined cohort (n = 13; **Figure 2C-E**).

The peak monoamine response level on a trial was defined as the maximum monoamine level within a 4-second window spanning 0 seconds before fractal cue onset to 4 seconds after fractal cue onset (i.e., window spanning fractal cue onset to 0.5 seconds post-liquid delivery; **Figure 2A**). We identified the time point of this maximum monoamine response level within the window of interest for each trial individually (**Figure 2D**) as well as for contiguous 2-trial averages (i.e., trials 1 and 2 were averaged together before finding the maximum monoamine level, and this process was repeated for trials 3 and 4, trials 5 and 6, and trials 7 and 8). The 2-trial averaging was used exclusively for visualization of the monoamine responses across conditioning. For the single-trial data, we regressed the timing of the peak monoamine response level against trial number for trials 2-8, as we observed that trial 1 engendered variable monoamine responses to both the cue and liquid reward that could arbitrarily anchor the trial regression at one or the other stimuli. This linear regression analysis was performed using the MATLAB function fitglm(). For the single-subject linear regression analyses, we took the individual-level beta coefficients to the second-level using a one-sample t-test to test for statistically significant patterns of monoamine response timings within each cohort (**Figure S3A**). We repeated this analysis for other features of the monoamine response besides the timing of the peak response level, namely the onset and offset of the positive monoamine cluster containing the peak response level (i.e., shaded regions beneath dopamine time series in **Figure 2C,D**). The onset and offset of the positive response cluster containing the maximum response level were found by identifying the samples at which the monoamine time series crossed zero at the start and end of the response cluster, respectively.

##### Ultimatum game data analysis and linear modeling

Similar to the analysis of the trace conditioning data, we investigated the adaptation in monoamine response timing across trials of the ultimatum game. We adopted a sliding-window approach to analysis (**Figure 3D**). For each participant, we first averaged together monoamine time series into overlapping 10-trial blocks with a 1-trial step size (i.e., block 1 is formed by averaging together trials 1-10; block 2 is formed by averaging trials 2-11; etc.), resulting in a Trials x Time matrix of size 51 x 61 (51 blocks by 61 monoamine samples; **Figure S5**). Then, for each ultimatum game cohort separately, we averaged together the block-level monoamine data for each participant within the cohort (e.g., the 13 participants in the High-to-High cohort were averaged together), yielding a single representative dataset for each cohort. We used this cohort-level representative dataset to visualize the temporal adaptations in peak monoamine response levels across time and across experimental variations (**Figure 3E,F**; **Figure S5,6**).

From these cohort-level representative datasets, we identified the time point of the maximum monoamine response level within a 3-second window of interest for each block. We focused our analyses on the period of an ultimatum game trial spanning the offer reveal to the choice selection screen, which are analogous to the trigger cue and response of a delay conditioning paradigm (**Figure 3C**). The 3-second window of interest was defined adaptively, with the window’s upper bound anchored to 1 second before the average response time for the given block. This upper bound was specified to avoid any putative action-related monoamine responses; additionally, if the average response time for a block was greater than 3 seconds, the window of interest was defined as 0-3 seconds post-offer reveal (i.e., window onset lower bound is the offer reveal and upper bound is 3 seconds post-offer reveal). In this way, the window of interest always spanned the period of a trial including the offer reveal and response screen onset. Furthermore, we observed that participants’ average response times decreased with experience, and the window of interest was designed to adapt proportionately to changes in response times so as to continue to avoid any action-related monoamine responses while also capturing any potential monoamine fluctuations preceding the offer reveal, such as in response to the partner proposing cue. The average response time for each block was determined separately for the combined High-to-High and High-to-Medium cohorts (i.e., “norm adaptation” groups; n = 28) and for the Choice Delay cohort (n = 16), given the differences in the trial response timing for the final 30 trials of the Choice Delay group. We assessed the reproducibility of the observed patterns in dopamine peak response timing by also performing this analysis using 3-, 5-, and 6-trial kernel widths for defining the sliding window (**Figure S5**).

To test whether the observed patterns in dopamine peak response timing for the High-to-High, High-to-Medium, and Choice Delay cohorts were specific to dopamine or whether the response patterns were also observed for serotonin and norepinephrine, we performed linear modeling of contiguous (i.e., non-overlapping) 10-trial blocks of monoamine data. In the sliding-window approach with 1-trial step size (supra), blocks 1, 11, 21, 31, 41, and 51 comprise non-overlapping, contiguous blocks of 10 trials spanning the entire experiment (i.e., trials 1-10, 11-20, 21-30, 31-40, 41-50, 51-60; **Figure S4**) and cleanly delineating the half-way point of the experiment where offer magnitude and choice timing are altered for the High-to-Medium and Choice Delay cohorts, respectively (i.e., contrasting trials 21-30 and trials 31-40).

Using these contiguous blocks of monoamine data, we defined cohort-specific linear models based on the observed dopamine peak response timings and fit the linear models to each cohort’s peak serotonin and norepinephrine response timings (**Figure S4**). The linear models were different for the three ultimatum game participant cohorts and were designed to capture the observed changes in dopamine response timings across blocks that appeared specific to each ultimatum game task variation (i.e., those observed in **Figure 3E,F**). Specifically, across blocks 1-3 (Trials 1-30), the linear models for all groups specify a negative change in dopamine peak response timing from block 1 (Trials 1-10) to block 2 (Trials 11-20) but no further change from block 2 to block 3 (Trials 21-30). Across blocks 4-6 (Trials 31-60), for all cohorts there is a monotonic backwards-shifting change in the peak DA response timing: for the High-to-High group, this shifting proceeds from around the offer reveal screen onset at block 4 to the onset of the partner proposing cue by block 6; for the High-to-Medium cohort, the backwards-shifting response begins at the choice selection screen onset (i.e., 1s post-offer) at block 4 and proceeds to shift to align to the offer reveal screen by block 5 and ultimately to align to before the offer reveal screen by block 6; and for the Choice Delay cohort, the backwards-shifting response initializes further back at the newly delayed response selection onset of 2.5s at block 4, before shifting backwards within the phase of a trial to align to the offer reveal by block 5, and then aligning to before the offer reveal by block 6.

To form linear contrasts that capture these cohort-specific patterns in the timing of the peak dopamine response within the phase of a trial, we segmented time within a trial into a set of temporally-ordered states: state 4 corresponds to the onset of the partner proposing cue, state 3 corresponds to the time prior to the onset of the offer reveal screen, state 2 corresponds to the onset of the offer reveal, state 1 corresponds to the onset of the response screen, and state 0 corresponds to the delayed response screen onset of the Choice Delay cohort. Note that the states were numbered in reverse order of their occurrence within a trial of the ultimatum game (i.e., state 4 is the earliest state of the trial, and state 1 or 0 is the final state of a trial), which was chosen to model the backwards movement of the dopamine response within the phase of a trial (i.e., dopamine responses move from state 1 backwards to state 2 over time, and then to state 3, etc.). Using this state indexing scheme, the linear model contrasts for each group were defined as: High-to-High model contrast = [1,2,2,2,3,4]; High-to-Medium model contrast = [1,2,2,1,2,3]; Choice Delay model contrast = [1,2,2,0,2,3].

---

**Figure S1.**
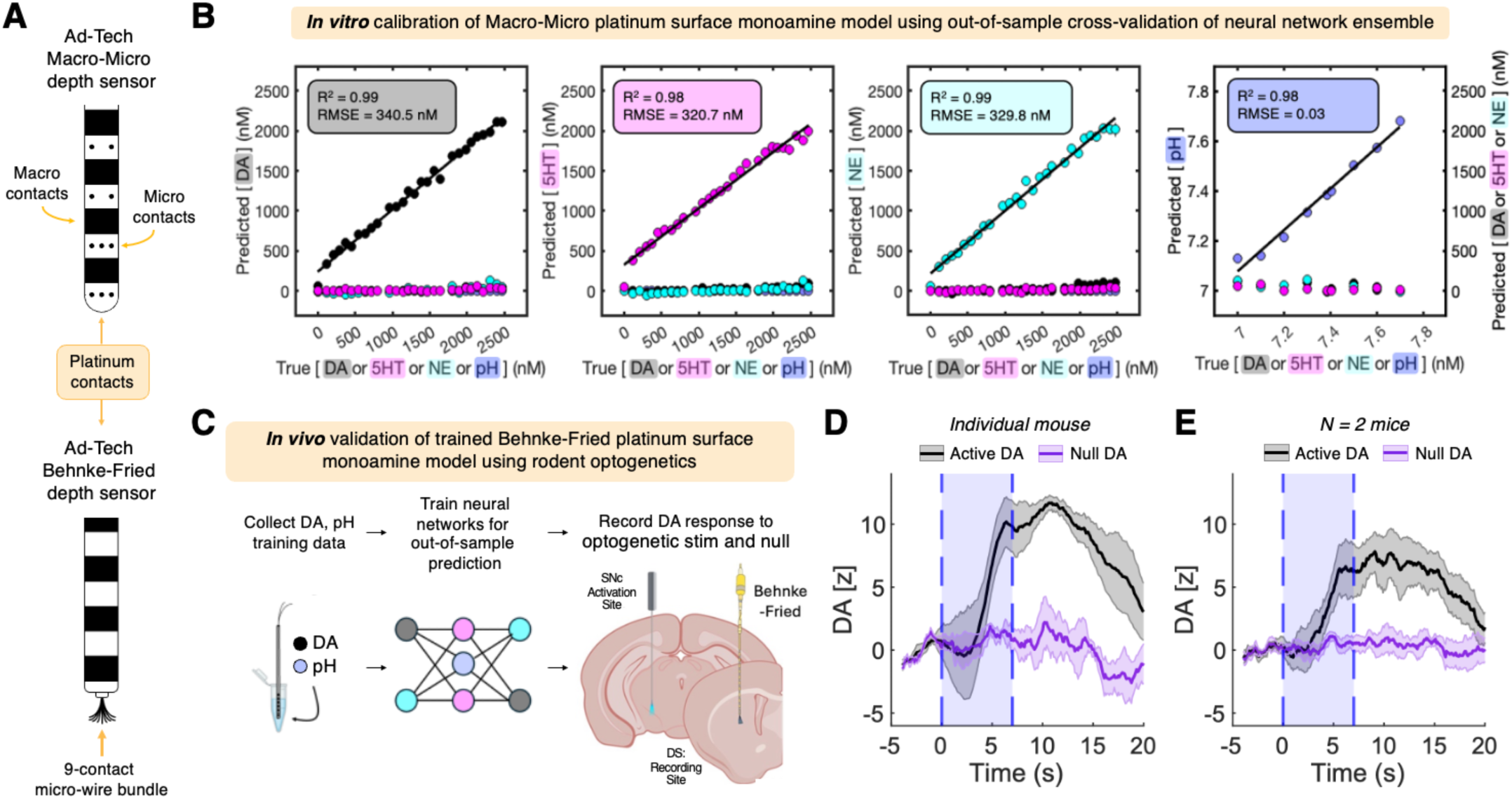
*In vitro* validation of human nasal sensor platinum surface and *in vivo* validation of platinum surface-based neuromodulator model training using rodent optogenetics. **(A)** Cartoon depicting the Ad-Tech Macro-Micro sEEG depth sensor used in the nasal recording sessions (top) and the Ad-Tech Behnke-Fried depth sensor used in the rodent optogenetic validation experiments. Note the location of the Macro-Micro sensor’s macro- and micro-wire contacts and the Behnke-Fried senor’s micro-wire bundle, which are all similar platinum contact surfaces. (B) Estimates of out-of-sample monoamine concentrations and pH levels from measured electrical currents. In each panel, the network-predicted monoamine concentration (in nM) is plotted against the true concentration. R^2^: r-squared. RMSE: root mean square error. These summaries represent a composite over all the probes included in this paper (N = 46 sensors). Data are plotted as mean ± SEM across sensors. **(C)** Illustration of sites for optogenetic stimulation (substantia nigra pars compacta, SNc) and electrochemical recording (dorsal striatum, DS). **(D)** Averaged dopamine (DA) time series aligned to the onset of a 7 second optical stimulation (shaded blue box) for trials using active optical stimulation (black trace; n = 3 trials) and inactive optical stimulation (purple trace; n = 3 trials) in a single mouse. The bold black and purple traces indicate the mean across trials of a condition; the surrounding shaded regions represents +/- 1 standard error of the mean (SEM). Note the increase in DA level in response to active optical stimulation, which is absent for trials using the null light outside the excitation spectra of ChR2; active and inactive optical stimulation trials were interleaved for all rodent optogenetic experiments. **(E)** Same as **(D)** but for averaged DA time series across two mice (including mouse shown in **(D**)) for active optical stimulation (black traces, n = 6 trials) and inactive optical stimulation (purple traces, n = 6 trials). Note that both mice contributed three active and inactive optical stimulations each.

**Figure S2.**
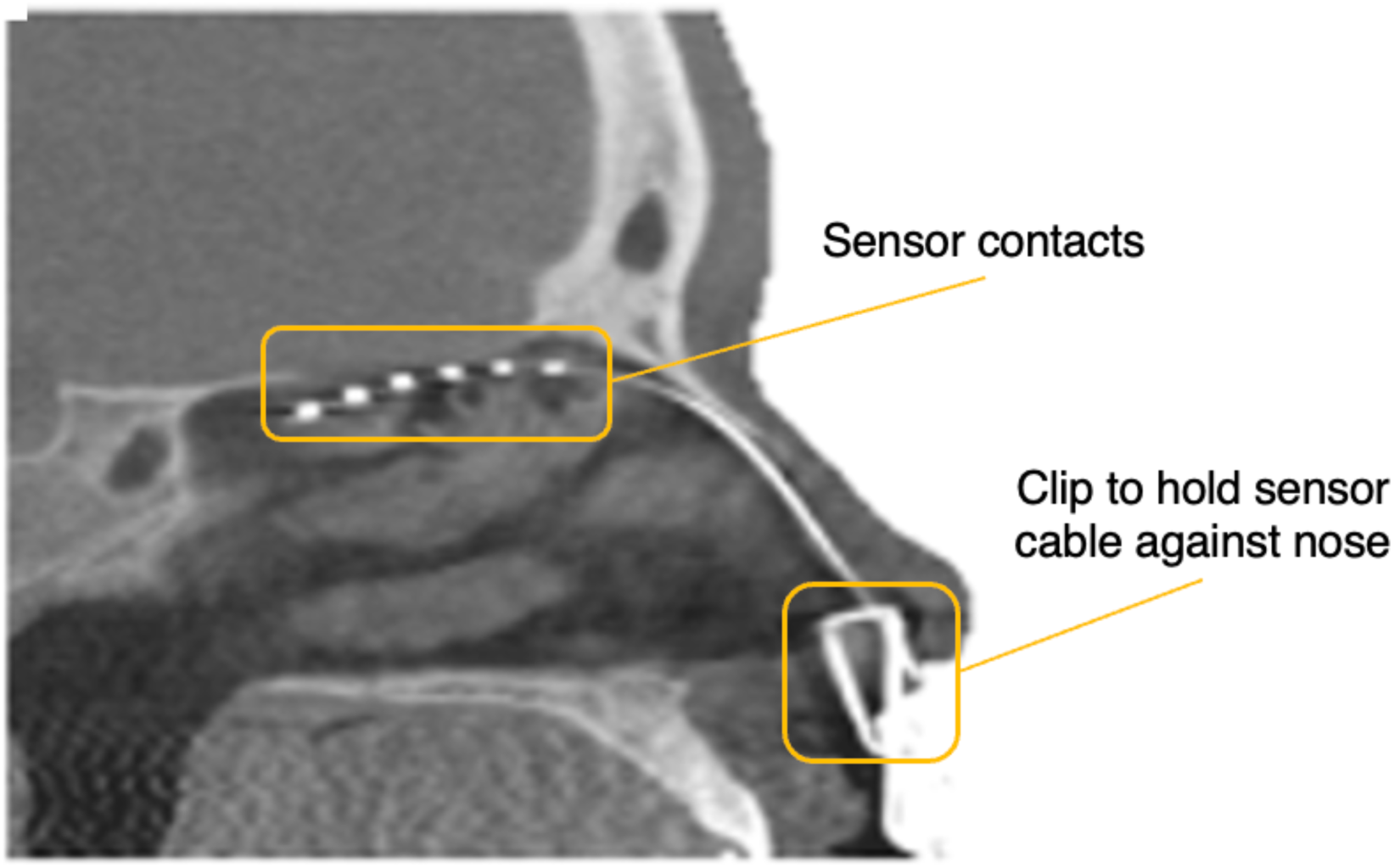
Head CT scan of sensor placement in nasal cavity. Note the sensor is placed the along the superior surface of the nasal cavity, just below the cribriform plate.

**Figure S3.**
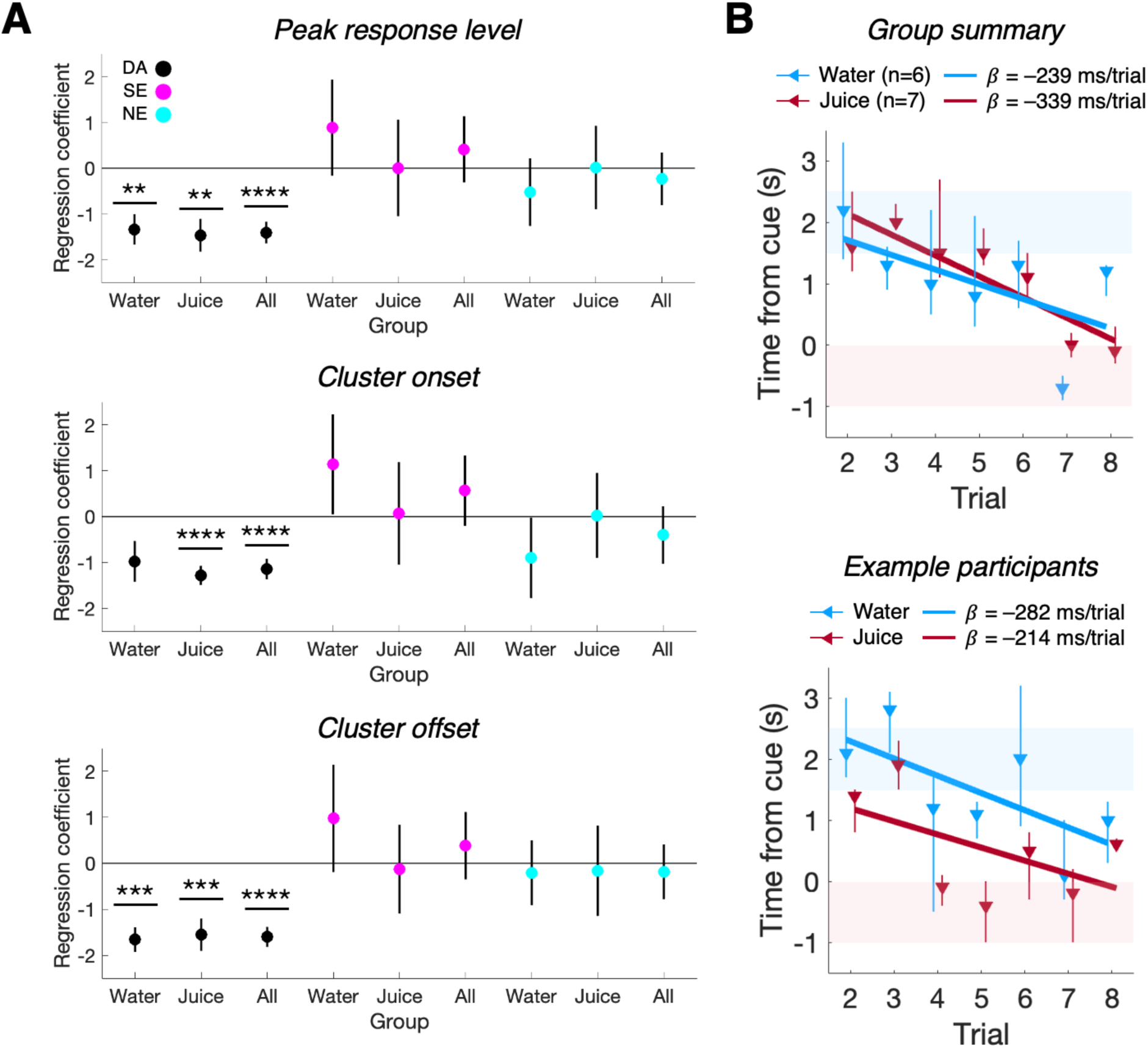
Regression analysis of monoamine response timing across conditioning trials. **(A)** Regression coefficients from linear regression of dopamine (DA; black), serotonin (SE; magenta), and norepinephrine (NE; cyan) response features with conditioning trial number. Colored circles indicate the group-level mean coefficient values, and vertical black lines represent one standard error of the mean (SEM). **p<0.01, ***p<0.005, ****p<0.001 one-sample t-test. Note that the regression coefficients for the DA (black circles) response peak timing across trials (top) were significantly negative for the Water cohort (t(5) = -4.05, p = 0.0098), the Juice cohort (t(6) = -4.10, p = 0.0064), and the combined cohort (t(12) = -5.96, p = 6.6e-5), and the regression coefficients for the DA response cluster onset (middle) and offset (bottom) timing across trials were similarly negative for the Water cohort (cluster onset: t(5) = -2.21, p = 0.078; cluster offset: t(5) = - 6.17, p = 0.002), the Juice cohort (cluster onset: t(6) = -6.15, p = 8.5e-4; cluster offset: t(6) = -4.12, p = 0.0045), and the combined cohort (cluster onset: t(12) = -5.04, p = 2.9e-4; cluster offset: t(12) = -7.36, p = 8.7e-6); comparatively, SE and NE response features were all not significantly different from zero. There were no differences between Water and Juice cohort regression coefficients for all DA, SE, and NE response features. **(B)** Regression of maximum DA time point on trial number for group-averaged (top) and single-subject (bottom) DA time series. Colored triangles indicate the timing of the maximum DA response, and vertical lines represent the onset and offset of the cluster containing the maximum DA response. Bold overlaid lines represent the slope (regression coefficient, *β*) of the group-level or subject-level DA peak timing shift across trials.

**Figure S4.**
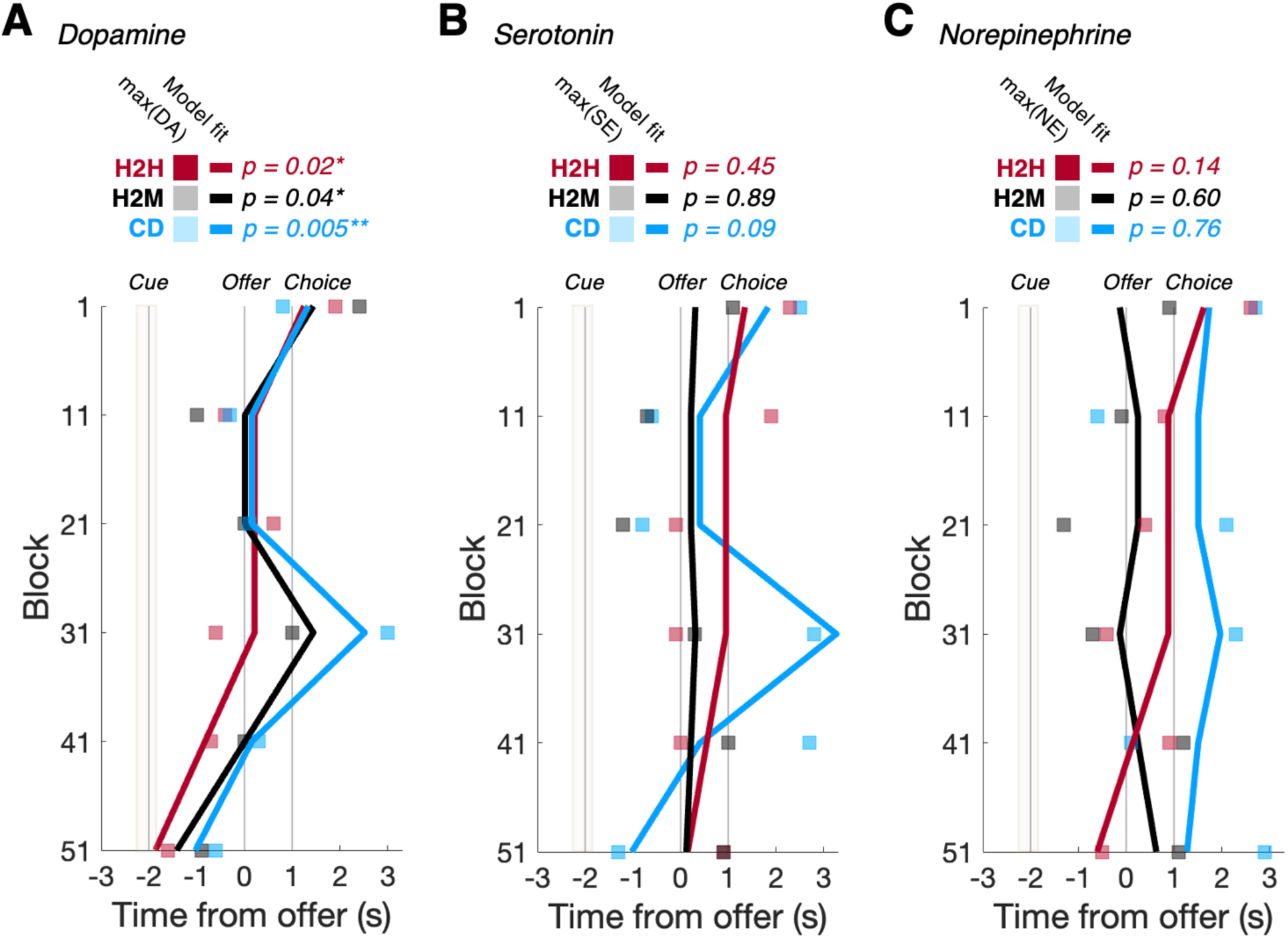
Linear modeling of monoamine response timing adaptation during the ultimatum game. **(A)** Linear modeling of dopamine (DA) peak response timing across contiguous 10-trial blocks of the ultimatum game for the High-to-High (red, n=10), High-to-Medium (black, n=15), and Choice Delay (blue, n=16) cohorts. Note the linear models (bold colored lines) fit to each group’s DA peak response timings (colored squares) were designed to capture the observed changes in group-specific DA response timings across blocks of each ultimatum game task variation. *p<0.05, **p<0.01. **(B)** and **(C)** same as **(A)** but for serotonin **(B)** and norepinephrine **(C)**. Note that the linear models that capture dopamine peak response timings in **(A)** do not account for observed peak response timings of serotonin **(B)** and norepinephrine **(C)**.

**Figure S5.**
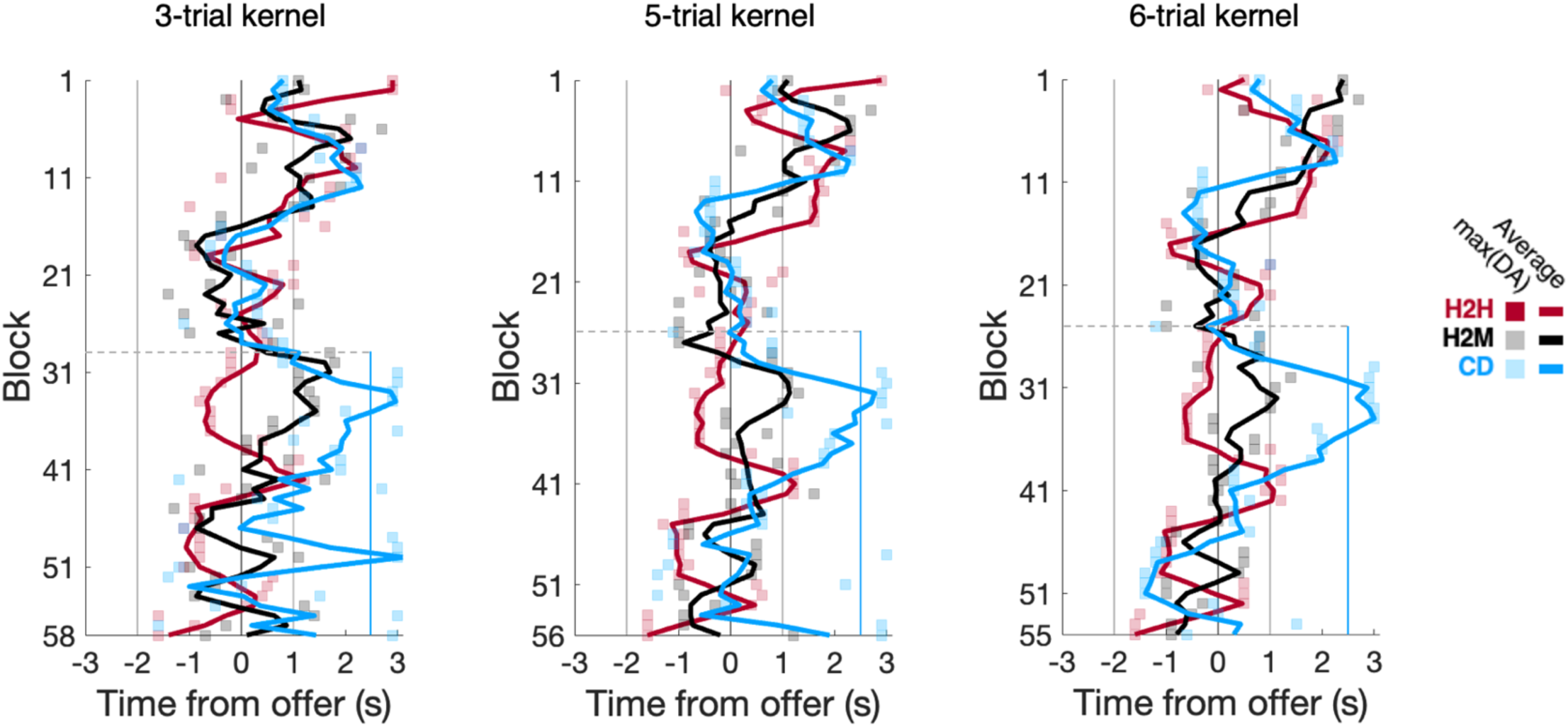
Sliding-window analysis of dopamine response shifting during ultimatum game with varying trial kernel widths. Timing of the peak dopamine (DA) responses (colored squares) for 3-trial (right), 5-trial (middle), and 6-trial blocks (right) of the UG task for the (left) High-to-High, (middle) High-to-Medium, and (right) Choice Delay cohorts. Bold colored lines indicate the simple moving average of the peak DA response over 3 blocks; vertical grey and blue lines indicate the timing of trial stimuli; shaded tan region centered on cue onset indicates the temporal jitter in the timing between cue presentation and offer reveal. Horizontal dashed lines show the transition in offer magnitude (H2M) or choice timing (CD), or the lack thereof (H2H), within the UG task version. Note the difference in total block number for each trial kernel width.

**Figure S6.**
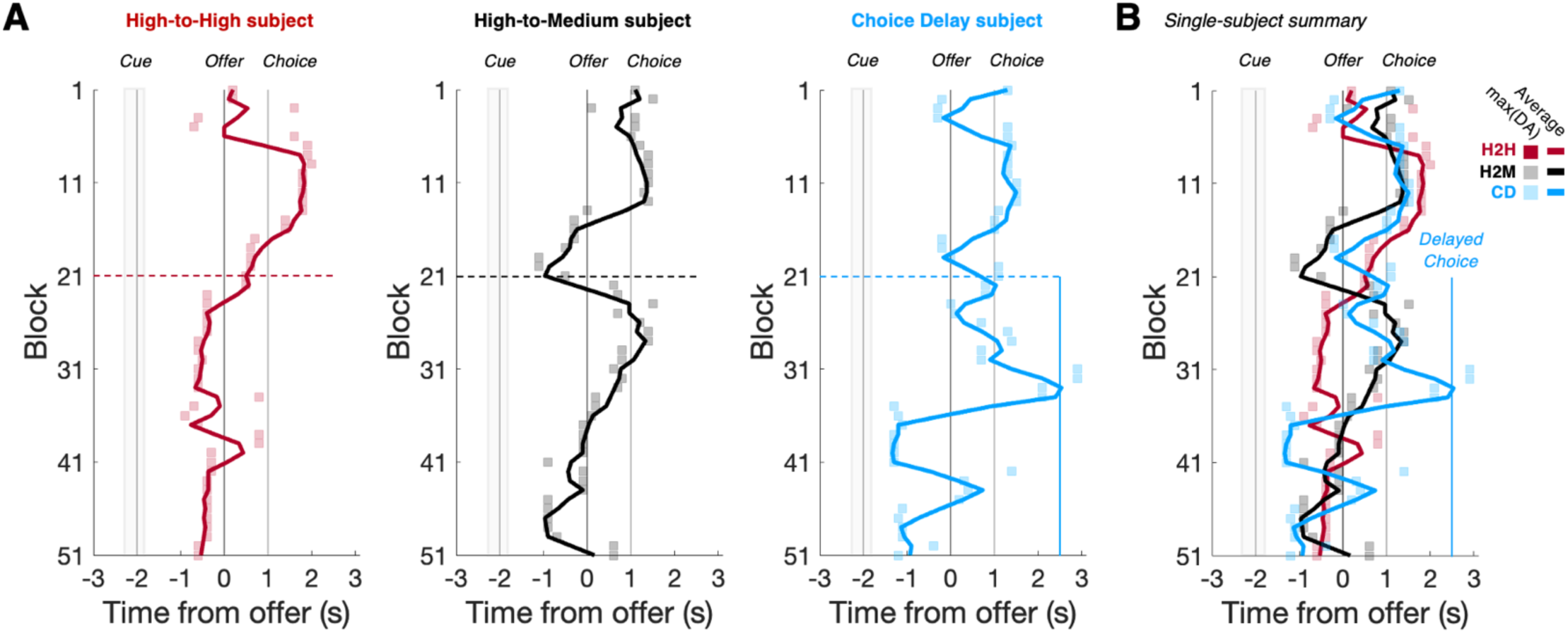
Single-subject dopamine response shifting during ultimatum game. **(A)** Timing of the peak dopamine (DA) responses (colored squares) for 10-trial blocks of the UG task for example single subjects from the (left) High-to-High, (middle) High-to-Medium, and (right) Choice Delay cohorts. Bold colored lines indicate the simple moving average of the peak DA response over 3 blocks; vertical grey and blue lines indicate the timing of trial stimuli; shaded tan region centered on cue onset indicates the temporal jitter in the timing between cue presentation and offer reveal. Horizontal dashed colored lines show the transition in offer magnitude (H2M) or choice timing (CD), or the lack thereof (H2H), within the UG task version. **(B)** Single-subject results from **(B)** overlaid.

**Table S1.** Participant demographics and visit information. This table displays the information of participants across sites, tasks, and conditions. Abbreviations: NWM, Northwestern School of Medicine; VT, Virginia Tech; UG, ultimatum game. * = subjects contributing datasets from multiple visits.

| Dataset | Session | Subject | Site | Task | Condition | Age | Sex | Nostril | Visit |
| --- | --- | --- | --- | --- | --- | --- | --- | --- | --- |
| 1 | 1 | 1 | NWM | UG | High-to-Medium | 46 | M | L | 2 |
| 2 | 2 | 2 | NWM | UG | High-to-Medium | 34 | F | R | 1 |
| 3 | 3 | 3 | NWM | UG | High-to-Medium | 32 | F | L | 1 |
| 4 | 4 | 4 | NWM | UG | High-to-Medium | 24 | M | L | 1 |
| 5 | 5 | 5 | NWM | UG | High-to-Medium | 42 | F | L | 1 |
| 6 | 6 | 6 | NWM | UG | High-to-Medium | 24 | M | R | 1 |
| 7 | 7 | 7 | NWM | UG | High-to-High | 19 | M | R | 1 |
| 8 | 8 | 8 | NWM | UG | High-to-High | 54 | M | R | 1 |
| 9 | 9 | 9 | VT | UG | High-to-Medium | 45 | M | L | 1 |
| 10 | 10 | 10 | VT | UG | High-to-Medium | 30 | M | R | 2 |
| 11 | 11 | 11 | VT | UG | High-to-Medium | 22 | F | L | 1 |
| 12 | 12 | 12 | VT | UG | High-to-Medium | 28 | F | R | 1 |
| 13 | 13 | 13 | VT | UG | High-to-High | 50 | F | L | 1 |
| 14 | 14 | 14 | VT | UG | High-to-High | 26 | M | L | 1 |
| 15 | 15 | 15 | VT | UG | High-to-High | 46 | M | L | 1 |
| 16 | 16 | 16 | VT | UG | High-to-High | 47 | F | R | 1 |
| 17 |  |  |  | Conditioning | Water |  |  |  |  |
| 18 | 17 | 17 | VT | UG | High-to-High | 35 | M | L | 1 |
| 19 | 18 | 18 | VT | UG | High-to-High | 34 | F | R | 1 |
| 20 | 19 | 19 | VT | UG | High-to-High | 23 | M | L | 1 |
| 21 |  |  |  | Conditioning | Water |  |  |  |  |
| 22 | 20 | 20 | VT | Conditioning | Water | 37 | F | R | 1 |
| 23 | 21 | 21 | VT | UG | High-to-High | 46 | F | L | 1 |
| 24 |  |  |  | Conditioning | Water |  |  |  |  |
| 25 | 22 | 22 | VT | UG | High-to-High | 46 | F | L | 1 |
| 26 |  |  |  | Conditioning | Water |  |  |  |  |
| 27 | 23 | 15* | VT | UG | High-to-Medium | 49 | M | L | 2 |
| 28 |  |  |  | Conditioning | Water |  |  |  |  |
| 29 | 24 | 16* | VT | UG | High-to-Medium | 47 | F | R | 2 |
| 30 | 25 | 23 | VT | UG | Choice Delay | 52 | M | R | 1 |
| 31 | 26 | 24 | VT | UG | Choice Delay | 23 | F | L | 1 |
| 32 |  |  |  | Conditioning | Juice |  |  |  |  |
| 33 | 27 | 25 | VT | UG | Choice Delay | 26 | F | L | 1 |
| 34 | 28 | 26 | VT | UG | Choice Delay | 22 | F | R | 1 |
| 35 | 29 | 27 | VT | UG | Choice Delay | 28 | F | R | 1 |
| 36 | 30 | 21* | VT | UG | Choice Delay | 46 | F | L | 2 |
| 37 |  |  |  | Conditioning | Juice |  |  |  |  |
| 38 | 31 | 28 | VT | UG | Choice Delay | 27 | M | R | 1 |
| 39 |  |  |  | Conditioning | Juice |  |  |  |  |
| 40 | 32 | 29 | VT | UG | Choice Delay | 29 | M | R | 1 |
| 41 |  |  |  | Conditioning | Juice |  |  |  |  |
| 42 | 33 | 30 | VT | UG | Choice Delay | 26 | F | L | 1 |
| 43 |  |  |  | Conditioning | Juice |  |  |  |  |
| 44 | 34 | 31 | VT | UG | Choice Delay | 31 | M | L | 2 |
| 45 | 35 | 10* | VT | UG | Choice Delay | 31 | M | R | 3 |
| 46 | 36 | 32 | VT | UG | Choice Delay | 39 | F | L | 1 |
| 47 | 37 | 33 | VT | UG | Choice Delay | 43 | F | R | 1 |
| 48 | 38 | 34 | VT | UG | Choice Delay | 23 | F | L | 1 |
| 49 |  |  |  | Conditioning | Juice |  |  |  |  |
| 50 | 39 | 35 | VT | UG | Choice Delay | 42 | M | L | 1 |
| 51 | 40 | 36 | VT | UG | Choice Delay | 46 | F | L | 1 |
| 52 | 41 | 37 | VT | UG | High-to-Medium | 75 | F | L | 1 |
| 53 | 42 | 38 | VT | UG | High-to-Medium | 49 | F | R | 1 |
| 54 | 43 | 39 | VT | Conditioning | Juice | 21 | F | L | 1 |
| 55 | 44 | 40 | VT | UG | High-to-Medium | 64 | F | L | 1 |
| 56 | 45 | 41 | VT | UG | High-to-Medium | 22 | F | R | 1 |
| 57 | 46 | 42 | VT | UG | High-to-Medium | 20 | F | L | 1 |

